# The Burden of Poor Household Drinking-Water Quality on HIV/AIDS Infected Individuals in Rural Communities of Ugu District Municipality, Kwazulu-Natal Province, South Africa

**DOI:** 10.1101/711002

**Authors:** C.M.N Khabo-Mmekoa, M.N.B Momba

## Abstract

This aim of this study was to ascertain whether household container-stored drinking water might play a role in the diarrhoeic conditions of HIV/AIDS patients and non-HIV-infected individuals of the rural communities who attended the Ugu District Municipal hospitals. Water samples were collected from the standpipes and household containers, and stool specimens were obtained from HIV/AIDS-positive and non-HIV/AIDS patients with diarrhoea. Significant correlations were established between the incidence of potentially pathogenic bacteria isolated from chlorinated household-stored water, and in stool specimens of HIV-positive patients with diarrhoea (r = P < 0.05). A combination of molecular analysis targeting the 16S rRNA gene and the restriction fragment length polymorphism and sequence analysis of the amplified gene for differentiating between species and strains of the bacterial pathogens was also applied to isolates obtained from stored-water samples and stool specimens. Similar sequences of *Klebsiella* spp., *K. pneumoniae, Escherichia coli, E. coli* O55: H7, *Proteus mirabilis*, and *Shigella boydii* were identified in both stored water and stools of HIV/AIDS-positive patients with diarrhoea. With the exception of *Proteus mirabilis*, none of these pathogens were identified in stool specimens of non-HIV-infected individuals with diarrhoea.

## 1. Introduction

The water service authority is tasked with the responsibility of improving the health of the general population by providing access to safe water. Lack of access to water and poor water quality may be as detrimental as not having water at all. Failure to provide safe drinking water poses a major threat to human health, while adequate water supply is a fundamental component of primary health. South Africa is classified as a water-scarce country and the supply of water does not meet the population’s growth [1]. However, the Constitution of the Republic of South Africa [2] states that every citizen has the right to have access to sufficient clean, safe drinking water. It should be a priority to improve and develop water infrastructure to ensure that communities are supplied with potable water.

Prior to 1994, 30 to 40% of South Africa’s population, which translates to an estimated 14 to 18 million people, had no access to clean or safe water [1;3]. By 2004, about 10 million people had been supplied with drinking water, thereby reducing the backlog that existed in 1994 [4]. By 2008, the estimated number of people who did not have a reliable source of drinking water had been lowered to six million [5]. In 2015, 3.64 million people in South Africa still had no access to an improved water supply [6]. Although this shows a fair commitment by the country to improve access to safe drinking water to all, studies have indicated that the focus has been mostly on metropolitan areas where the infrastructure for water treatment and supply to consumers is of a high quality compared to rural areas where it is poor or non-existent [7-12].

The implementation of communal standpipes and storage of treated drinking water in the dwellings remains the main mode of drinking water supply in most rural communities of South Africa. However, the provision of treated drinking water through this mode of access is not appropriate to maintain the safety of drinking water. Previous studies conducted on container-stored water worldwide have indicated the deterioration of the microbiological and physicochemical quality of this water source between the time of collection and storage [13-16].

Some studies have indicated that long storage times and the types of the container affect the microbiological and physicochemical quality of water collected and stored for drinking purposes by households [8; 13]. During storage, the quality of the water deteriorates, allowing the regrowth and survival of pathogenic microorganisms on the surfaces of household containers [8]. Other studies have attributed the deterioration of this water source to the living conditions and hygiene practices [17-20].

Unsanitary methods for dispensing water from household storage vessels, including contaminated hands and dippers and inadequate cleaning of vessels, number of children and socio-cultural status, have been shown to lead to the accumulation of sediments and pathogens [21; 22]. Even piped water supplies of adequate microbiological quality can pose a risk of infectious disease if they become contaminated due to unsanitary collection and storage conditions and practices within households [23]. Clearly, container-stored water is a critical public health concern.

Exposure to poor water quality could contribute to the spread of diseases such as cholera, typhoid fever, diarrhoea and liver problems [24; 25]. The impact of water-borne disease is significant. The World Health Organization [26] has estimated that about 88% of diarrhoeal diseases in the world are attributed to unsafe water and a lack of sanitation and hygiene. These diseases result in debilitating effects on rural communities, resulting in 3.1% of annual deaths recorded [9; 11; 27; 28].

In South Africa, diarrhoea is among the top ten causes of death, claiming 13 000 lives annually [29; 30]. The microbiological safety of drinking water is, therefore, a critical matter, especially for immuno-compromised people such as HIV/AIDS individuals. Due to their weakened immune system, diarrhoeal diseases transmitted by waterborne agents contribute significantly to their morbidity and mortality [31]. HIV-infected individuals have a greater need for potable water than uninfected individuals [32].

In South Africa, a few studies showing the link between the presence of some diarrhoea-causing bacteria pathogens in the stools of HIV-positive individuals and those found in their household drinking water have been conducted in rural areas of the Limpopo Province [33] and the Eastern Cape Province [34]. However, none of these studies have linked the deterioration of the quality of household-stored water to that of drinking water supplied through standpipe systems after treatment. Moreover, all these studies were based on a combination of culture-based methods and conventional PCR amplification of species-specific genes of pathogenic bacteria. More molecular techniques such as restriction fragment length polymorphism (RFLP) and sequencing are therefore required to establish whether the incidence of pathogenic bacteria in household container-stored drinking water plays a crucial role in the diarrhoeic conditions of immunocompromised people, such as HIV/AIDS patients residing in rural areas of underdeveloped countries in general and South Africa in particular. The most important fact is that to date no investigation of this nature has been conducted in the KwaZulu-Natal Province, while this province has the highest prevalence rate (39.5%) of HIV-infected individuals in South Africa [35].

The present study investigated whether the quality of drinking water at the point of supply (standpipe) was the first source for the deterioration of container-stored water quality, which might play a potential role in the diarrhoeic conditions of HIV/AIDS-positive and non-HIV/AIDS patients living in the Ugu District Municipality in the KwaZulu-Natal Province of South Africa. Conventional and molecular methods such as PCR targeting the 16S rRNA gene, PCR restriction fragment length polymorphism (RFLP), and sequence analysis of 16S rRNA amplified genes were used for the isolation, characterisation, and identification of pathogenic bacteria from standpipe water, water stored in households and stools of HIV/AIDS-positive and non-HIV/AIDS patients attending the Murchison and Port Shepstone hospitals.

## 2. Materials and Methods

### 2.1 Description of the study site, water supply, and storage

This study was carried out in the Ugu District Municipality in KwaZulu-Natal for a period of 11 alternating months between November 2008 and November 2009 and in April 2015. This district was selected because of its high prevalence rate of HIV (44%), high unemployment rate (30%), the backlog regarding the provision of water services to the population (70%), and the population distribution by race (89% black, 5% white, 3% Asian 1% Coloured, 2% other) (Statistics – Ugu District Municipality, 2010). There are two main referral hospitals, namely the Murchison and Port Shepstone hospitals, which serve the Ugu District Municipality. Statistical data of HIV/AIDS patients were obtained from the clinics at the Murchison and Port Shepstone hospitals. This information assisted in identifying the locations where water sampling was subsequently done.

The Boboyi, Bomela and Gamalakhe areas receive their drinking water from the Boboyi Water Purification Plant (**S1: Fig.1**). This plant abstracts its intake water from the Umzimkhulu River and uses conventional methods to produce final drinking water. The treatment includes the following multiple barriers: coagulation, flocculation, sedimentation, rapid sand filtration, and chlorination. In these areas, the storage of the treated water is crucial as the communities receive their water from standpipe systems. Drinking water is kept in plastic containers in the home for a period of up to three days. This storage period was also taken into consideration to assess the deterioration of drinking water over time. It was recorded during the study period that water was continuously added to these containers (four or five times) without cleaning them. The scoop used to draw water from the containers was placed on top of the containers and it remained uncovered.

### 2.2 Ethics clearance

To conduct the present study, ethics clearance was obtained from the Tshwane University of Technology (TUT), the Provincial Department of Health in KwaZulu-Natal and the relevant hospital clinics. Informed consent to participate in the study was obtained from patients, mothers of babies or patient guardians who granted permission to collect the stool specimens of HIV/AIDS and non-HIV-infected individuals by the hospital nurses. For the household drinking water samples, the informed consent was obtained from the owners of houses. Prior to sample collections, a clear justification of the aim and objectives of the study was provided to the study participants.

### 2.3 Collection of drinking water samples

Based on statistical data obtained from the clinics at the Murchison and Port Shepstone hospitals, the sampling programme was performed only in the areas where the target patients could be found although their names remained unknown to the research team. A total of 27 standpipes, which supplied drinking water to the selected 220 houses were considered for this study. A sampling of the standpipe drinking water was done concomitantly with that of household containers, using internationally accepted techniques and principles. However, it is important to mention that water samples collected from household containers could have a storage period varying from one to three days. For the purpose of this part of the study, water samples were randomly selected: 24 samples from standpipes and 72 samples from household containers (24 from each site) of Boboyi, Bomela, and Gamalakhe. Our intention was just to obtain an initial opinion on the incidence of pathogenic bacteria in drinking water supplied to communities and to determine whether this water might be a potential source of pathogens in household container-stored water, which might play a role in the diarrhoeic conditions of the HIV patients visiting the clinics while living in the above-mentioned rural areas.

The concentration of the residual chlorine and the turbidity level in drinking water samples were determined onsite according to the procedures outlined in the Standard Methods [36]. Water samples were stored on ice and transported to the laboratory at the Boboyi Water Treatment Plant for initial microbiological analyses within 6 to 8 h of being collected.

### 2.4 Collection of stool specimens

One hundred and twenty diarrhoeal stool samples from confirmed HIV-positive patients and 44 diarrhoeal stool samples from HIV-negative patients visiting the clinics and residing in the target areas were collected by hospital clinicians They were collected in clean, sterile wide-mouthed bottles (Merck, Johannesburg, South Africa) and transported to the National Health Laboratory Services at the Port Shepstone hospital laboratory for initial bacterial analyses within 4 to 6 h of being collected. All the patients’ locations, their diarrhoeic conditions, and their status were recorded at the hospitals. Confidentiality on the HIV status of patients was maintained throughout the project.

### 2.5 Isolation and detection of faecal coliforms and pathogenic bacteria

Standard methods with some modifications were used for the isolation and detection of the targeted bacterial enteric pathogens (*Salmonella Typhimurium, Shigella dysenteriae*, and *Vibrio cholerae*) from water samples and stool specimens [37; 38]. The culture media consisted of Chromocult® coliform agar (CCA), tetrathionate broth (TTB), thiosulphate-citrate-bile salt-sucrose (TCBS) agar, xylose lysine deoxycholate (XLD) and alkaline peptone water. They were prepared according to the manufacturer’s instructions (Merck, South Africa) and Standard Methods [38].

Faecal coliforms were quantified by membrane filtration techniques using membrane faecal coliform (m-FC) agar and Chromocult® coliform agar (CCA), and plates were incubated at 44.5 °C for 24 h. For *Salmonella* and *Shigella* spp., 500 mL drinking water samples were filtered through the sterile 42 mm diameter membrane of 0.45 µm pore size (Millipore). After filtration, the membranes were aseptically immersed in 50 mL of sterile Selenite broth (Merck, South Africa), and aerobically incubated at 36 ± 1 °C for 18 h to 24 h. Thereafter, 1 mL of the pre-enriched suspensions for each drinking water sample was added to sterile tubes containing 9 mL of TTB (Merck, South Africa). The suspensions were subjected to incubation at 36 ± 1 °C overnight. The detection of presumptive *Salmonella* and *Shigella* spp. was performed by the streak-plate method using xylose lysine deoxycholate (XLD) agar (Merck, South Africa) and the plates were incubated at 36 ± 1 °C overnight. For *Vibrio* spp., similar techniques were used but the pre-enrichment of these species was done using 100 mL of double-strength alkaline peptone water (pH 8.5) and the incubation period was 6 h to 8 h at 36 ± 1 °C. The detection of *Vibrio cholerae* was performed using thiosulphate-citrate-bile salts-sucrose (TCBS) agar. The same procedure was used for stool samples with the exception that a pea-sized stool specimen was directly inoculated in the respective enrichment broths for detection of *Salmonella*, S*higella* and *Vibrio cholerae*. All the tests were done in triplicate for the isolation of the aforementioned presumptive organisms. For water samples and stool specimens, three to five individual colonies differing in size, form, and colour were randomly selected, transferred into the slant bottles containing the relevant selective media, incubated at 36 ± 1 °C for 24 h and taken to the Tshwane University of Technology Water Research Group laboratory for confirmation and molecular analysis.

### 2.6 Identification of bacterial strains from water samples and stool specimens

#### 2.6.1 Culture-based methods

The selected colonies were transferred onto their corresponding selective media by the streak plate method and incubated at 36 ± 1 °C for 24 h. The colonies were further purified by the same methods at least three times using nutrient agar (BioLab) before Gram-staining. Oxidase tests were then performed on those colonies that were Gram-negative. The 20E API kit was used for the oxidase-negative colonies and the strips were incubated at 36 ± 1 °C for 24 h. The strips were then analysed, and the strains identified using API LAB PLUS computer software (BioMérieux, Marcy-l’Étoile, France).

### 2.7 Molecular techniques

#### 2.7.1 Extraction of the total genomic DNA

Total genomic DNA of the selected colonies was extracted using the DNeasy® DNA purification kit (QIAGEN) and ZR Fungal/Bacterial DNA MiniPrep™ kit (ZYMO Research, USA) according to the manufacturer’s instructions. The DNA concentrations were determined using the NanoDrop™ 2000 spectrophotometer (Thermo Fisher Scientific). The integrity of the purified DNA template was assessed by conventional agarose gel (1% [wt/vol] electrophoresis (Bio-Rad), and the purified DNA samples were stored at 4 °C until further use.

### 2.8 PCR amplification of the 16S rRNA gene

Universal primers 27F [39] and 1507R [40] were used in the polymerase chain reaction (PCR) reactions for the amplification of the 16S rRNA gene of each of the isolates. The reaction mixtures used in the PCR steps contained 12.5 μL of DreamTaq Master mix (2x) (Fermentas GmbH, 140 St. Leon-Rot, Germany), 0.5 μL of each primer (10 pmol), 8.5 μL of nuclease-free water (Fermentas, 140 St. Leon-Rot, Germany) and 5 μL of template DNA. The PCR reaction mixtures were placed in an MJ MINI thermal cycler (Bio-Rad), and the following thermal cycling conditions were used: pre-denaturation for 10 min, followed by 35 amplification cycles of denaturation at 94 °C for 30 s, annealing of primers with template DNA at 55 °C for 30 s and primer extension at 72 °C for 30 s. This was followed by a final extension at 72 °C for 7 min. The PCR amplicons were resolved through electrophoresis of 1% (w/v) agarose gel stained with ethidium bromide, followed by visualisation under ultraviolet light. The FastRuler™ Low Range DNA ladder (Fermentas, 140 St. Leon-Rot, Germany) was included in all gels as a size marker. All results were captured using a gel documentation system (Syngene, Cambridge, UK). The PCR amplicons were subjected to restriction analysis.

### 2.9 Restriction analysis of PCR amplicons

In order to select representative isolates for sequencing, all PCR amplicons were subjected to restriction fragment length polymorphism (RFLP) analysis. For this purpose, 10 μL of the 16S rRNA amplicons were digested with *Taq1* and *Cs6pI* (Fermentas) according to the manufacturer’s instructions. The restriction digests were resolved through electrophoresis of conventional 1.5% (w/v) agarose gel stained with ethidium bromide, followed by visualisation under ultraviolet light. The HyperLadder™ 1 kb, 100 lanes (Bioline Products, South Africa) was included in all gels as a size marker. All results were captured using a gel documentation system (Syngene, Cambridge, UK). The restriction patterns were determined manually, and for every five similar profiles, one isolate was selected for sequencing.

### 2.10 Sanger sequencing of the 16S rRNA gene

After grouping the isolates using the PCR-RFLP, the genomic DNA from the representative water sample and stool specimen isolates were amplified using the existing 27 F and 1507 R primers as described above. The 1500 PCR amplicons were further studied by conventional Sanger (dideoxy) sequencing in both directions using 27 F and 1507 R primers. For this purpose, BigDye™ for ABI 3130XL was used according to the manufacturer’s instructions and the gel was run on the 3130XL sequencer. All the sequences were inspected and manually corrected using BioEdit v.5.0.9 [41]. For the identification, the National Centre for Biotechnology Information (NCBI; *http://www.ncbi.nlm.nih.gov/*) nucleotide database was used to compare the sequences of bacteria isolated from water and stools using BLASTn.

### 2.11 Statistical analysis

Data were analysed using the SPSS Statistics 17.0 program. Pearson’s correlation (r_p_) coefficient (r = rho) was used to test the significant difference between the incidence of microorganisms in water samples and the concentration of free chlorine residual, and also the turbidity level in stored water over time. The significant linear relationship was also established between the incidence of microorganisms in water and in stool specimens. The statistical significance level was set at p ≤ 0.01 and p ≤ 0.05. Spearman’s rank correlation coefficient (r_s_ or rho) was run concurrently to also establish statistical significance.

## 3. Results

### 3.1 Turbidity of drinking water

The average turbidity levels of drinking water supplied through standpipes to Bomela and Boboyi communities were found to range between 1 and 4.7 NTU. The highest turbidity level was recorded in > 50% of all the drinking water collected samples collected from Bomela and Gamalakhe and the lowest was recorded in < 50% of the drinking water samples collected in Boboyi. Out of the 30 selected standpipe drinking water samples, 27.3%, 36.4%, and 27.3% were found to have turbidity levels ranging between 1 and 1.7 NTU in Gamalakhe, Bomela, and Boboyi respectively (**S2: Fig. 2**).

Moreover, **S3: Fig. 3** depicts the turbidity levels of drinking water over the storage period in a household container. The average turbidity levels ranged between 1 and 8.1 NTU. Out of the 72 selected container-stored drinking water samples, results revealed that only 29%, 4%, and 9.1% were found to have turbidity levels ranging between 1 and 1.7 NTU in Gamalakhe, Bomela, and Boboyi, respectively. The highest turbidity values, ranging between 7.6 and 8.1 NTU, were found in 4.5% of container-stored water samples collected from Boboyi and Bomela (**S4: Fig. 4**). In general, there was a marked increase in turbidity values over the storage period in spite of the fluctuations that occurred from time to time. Statistical evidence showed a significant difference and a positive correlation between turbidity values in stored water and the storage period (r = 0.49, r = 0.05).

### 3.2 Chlorine residual in drinking water

The presence of a chlorine residual in drinking water is to inactivate bacteria and some viruses that can cause diarrhoeal disease and to protect the water from being re-contaminated. Other studies have reported on the growth of faecal coliforms and presumptive pathogenic bacteria in chlorinated water [13; 42; 43]. The concentrations of chlorine residual concentrations in all water samples collected from Bomela (0.51 mg/L), Gamalakhe (0.51 mg/L), and Boboyi (0.42 mg/L) were within the recommended limits (0.3 to 0.6 mg/L) [44]. (DWAF, DoH and WRC, 1998). The findings of this study have indicated that high turbidity levels affected the efficiency of free chlorine residual in inhibiting the growth of bacteria in household container-stored water (**S4: Fig.4**). The Boboyi standpipe had the lowest chlorine residual concentration range of 0.05 mg/L in water samples collected during the study period and the highest was 0.815 mg/L. The lowest chlorine residual concentration was 0.06 mg/L and the highest was in the range of 1.095 mg/L”) As to Gamalakhe, the lowest concentration was 0.12 mg/L of free chlorine residual was found in water samples and the highest in the range was 0.895 mg/L (Figure 3).

**Fig. 1:**
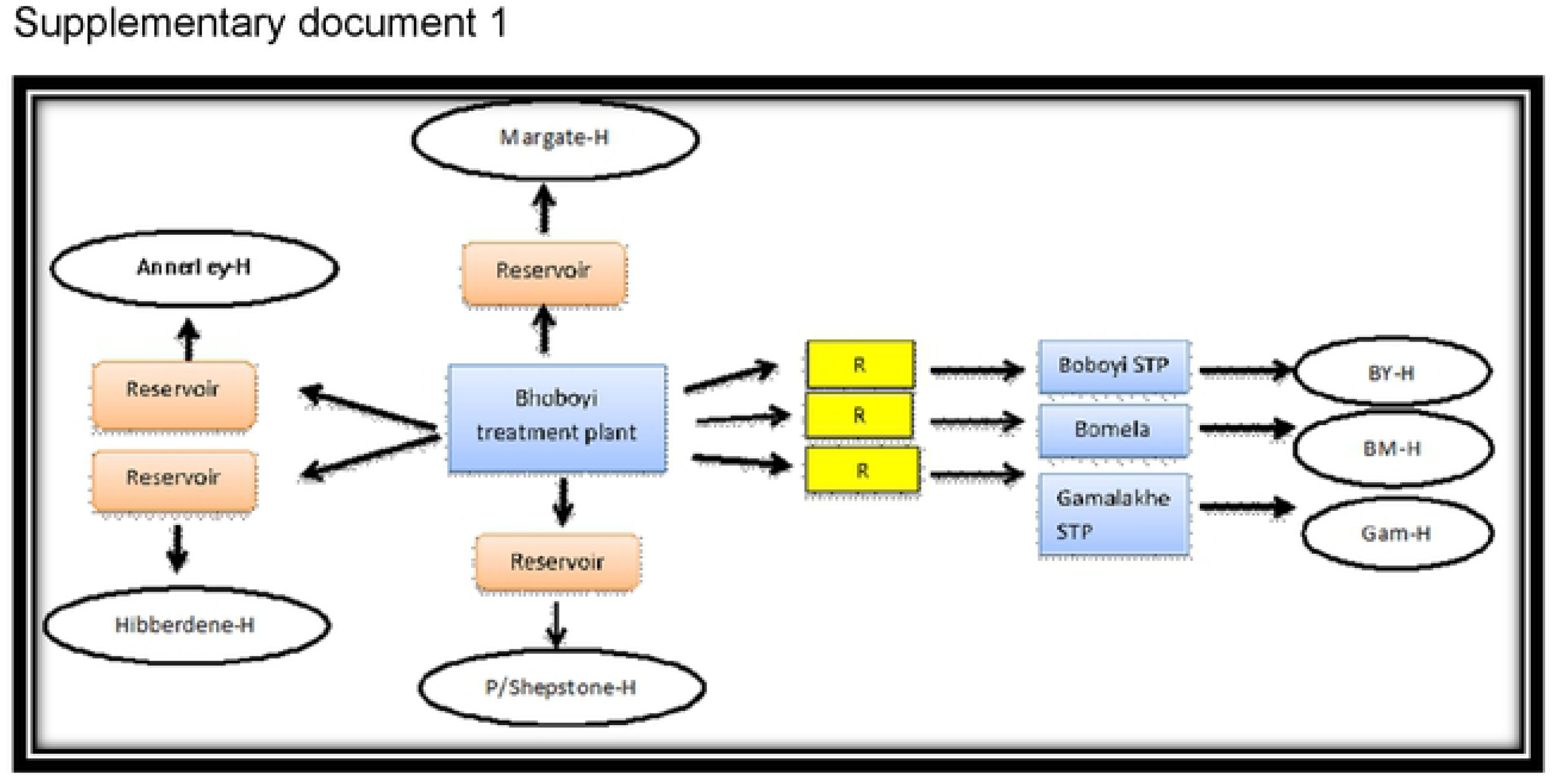
Mode of access to municipal drinking water supply in target urban and rural areas (H: house; R: reservoir; STP: Standpipe; BY-H: Boboyi house; BM H: Bomela house; Gam-H: Gamalakhe house)

**Fig. 3:**
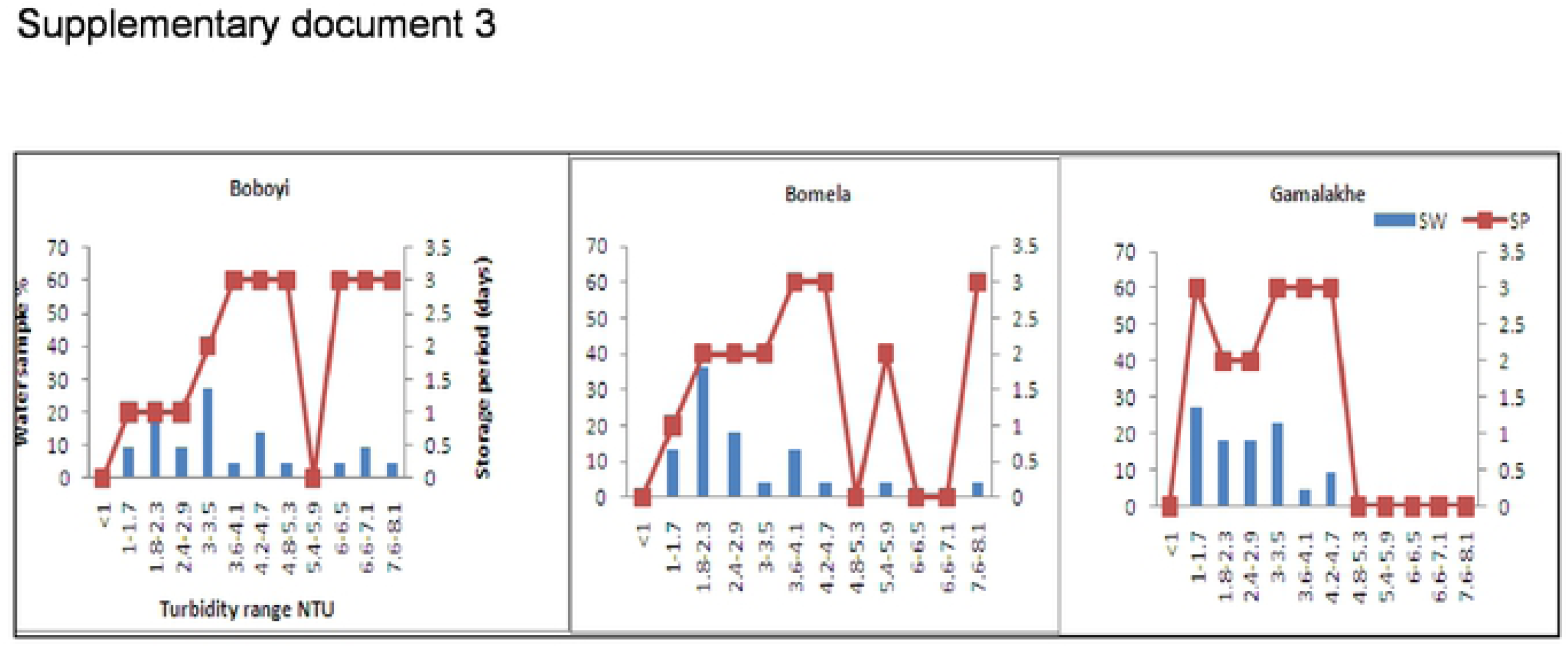
Increased turbidity(NTU) levels in stored household water samples(%) collected from Boboyi, Bomela and Gamalakhe over time (SP = storage period, SW = stored water)

**Fig. 4:**
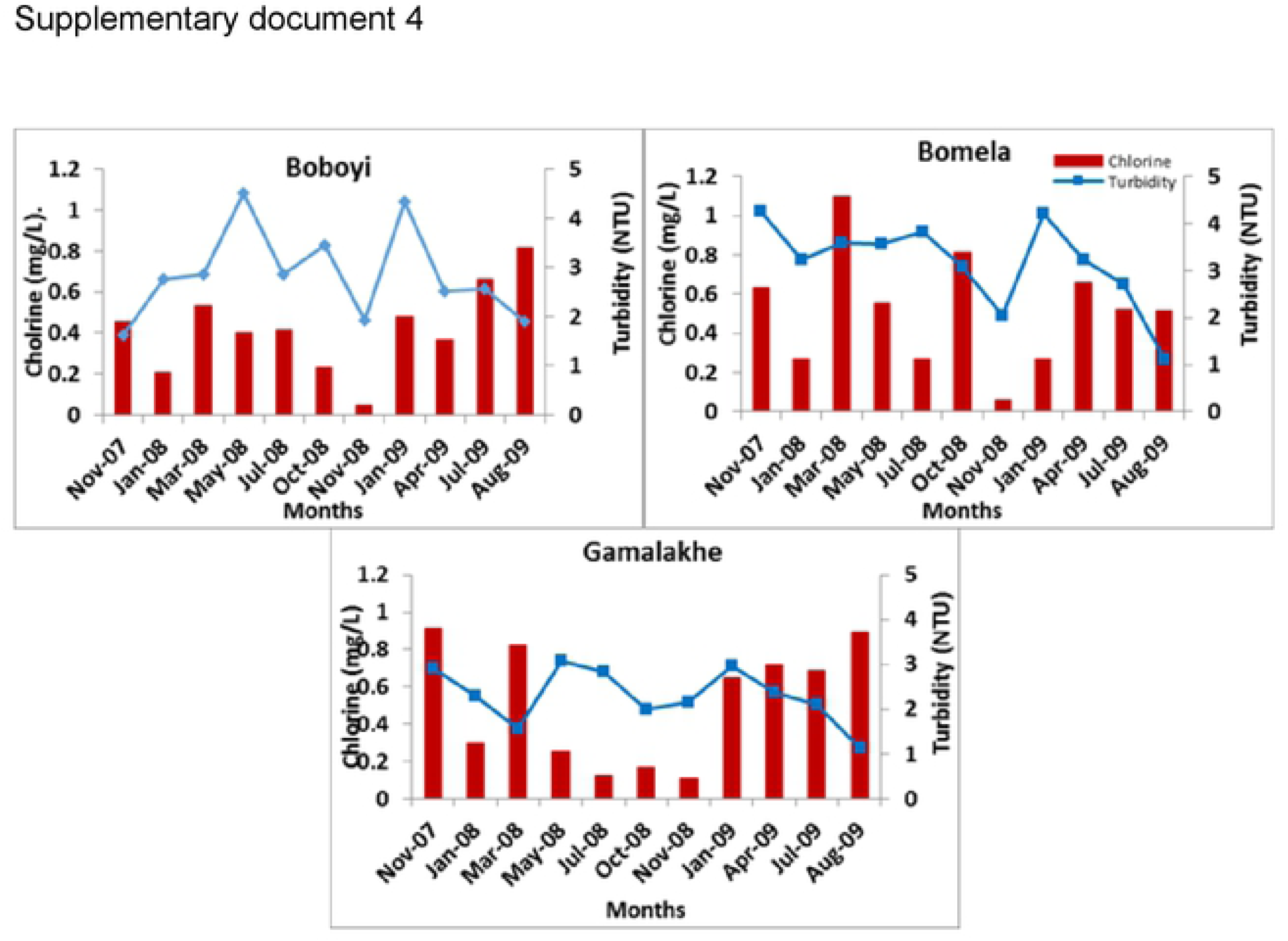
Average chlorine values (mg/L) in stored household container water samples collected from Boboyi, Bomela, and Gamalakhe

All container-stored drinking water samples collected from Boboyi were found to have chlorine residual concentrations ranging between < 0.05 mg/L and 0.8 to 0.9 mg/L. The majority of the water samples in this area had chlorine residual concentration ranges of 0.8 – 0.9 mg/L and 0.3 – 0.4 mg/L. For Bomela the chlorine residual concentrations ranged between < 0.05 mg/L and 1.5 mg/L, with 0.4 to 0.5 mg/L recorded for the largest number of samples (27.3%). None of the stored-water samples collected from Gamalakhe were found to have chlorine residual concentrations < 0.05 mg/L. The chlorine residuals ranged between 0.05 and 1.5 mg/L. An equal number of water samples (22.7%) were found to have between 0.1 and 0.2 mg/L and between 0.8 and 0.9 mg/L free chlorine residual. A similar percentage of water samples (13.6%) originating from Bomela and Gamalakhe container-stored water was found to have the highest chlorine residual concentration of 1.5 mg/L.

#### Microbiological characteristics of drinking water and stool specimens

None of the target bacteria were recorded in water samples collected from the standpipes. Using culture-based methods, almost a similar rate of 31.8% of water samples with faecal coliforms were recorded in Boboyi and Bomela household-stored water, while a low rate of 8.3% was observed in Gamalakhe. Presumptive *Salmonella* spp. was found in half (50%) of Bomela household-stored water and also in more than 33% of Gamalakhe and Boboyi household-stored water. Presumptive *Shigella* spp. was detected at rates of ≥ 50% in household containers of the three villages and presumptive *Vibrio* spp. was detected at a higher rate compared to the rates recorded for other organisms. Correlations were found between the high levels of turbidity in container-stored water and the incidence of presumptive bacterial pathogens (at 0.01 level: r_p_ = 0.52, p-value = 0.003; at 0.05 level: r_s_ = 0.43, p-value = 0.019 for presumptive *Shigella/Salmonella*; and at 0.01 level: r_s_ = 0.48; p-value = 0.007; at 0.05 level: r_p_ = 0.39, p-value = 0.034 for presumptive *Vibrio*).

There were significant correlations at the 0.05 level between the incidence of microorganisms in household-stored water samples and in stool specimens of HIV-positive patients with diarrhoea (r_s_ = 0.36; p-value = 0.048 for presumptive *Shigella/Salmonella*; and r_s_ = 0.37; p-value = 0.047 for presumptive *Vibrio*).

A combination of PCR targeting 16S rRNA gene, PCR-restriction fragment length polymorphism (RFLP), and sequence analysis of 16S rRNA amplified genes resulted in the identification of similar sequences of specific bacterial pathogens from household container-stored water and those from stool specimens of HIV/AIDS-positive patients with diarrhoea. Table 4.1 depicts the pathogens detected in both stored water samples and stool specimens. With the exception of *Shigella*, none of the target presumptive pathogenic bacteria were identified using a combination of molecular techniques. These identified bacteria included *Klebsiella* spp., K. *pneumoniae, E. coli, E. coli O55: H7*, and *Shigella boydii* (Table 4.1). They were detected in both the selected household container-stored drinking water samples and stool specimens of HIV-positive individuals who suffered from diarrhoea during the study period. With the exception of *Proteus mirabilis*, none of these pathogenic bacteria were found in the stool specimens of HIV-negative patients who visited the hospitals for diarrhoea. *Proteus mirabilis* was identified in the stool specimens of HIV-negative patients who lived in Gamalakhe village.

## 4. Discussion

The survival and growth of microorganisms in the water is dependent on several factors that include the temperature of the water, the turbidity, level of chlorine, pH, and other sources for their nutrients. The highest turbidity level was recorded in > 50% of all the drinking water samples collected from Bomela and Gamalakhe the lowest was recorded in Boboyi drinking water (< 50%). Out of the 30 selected standpipe drinking water samples, 27.3%, 36.4%, and 27.3% were found to have turbidity levels ranging between 1 and 3.5 NTU in Gamalakhe, Bomela, and Boboyi, respectively (**S2: Fig.2**). The turbidity levels recorded for similar villages regarding stored water samples were 29%, 4% and 9.1% of, respectively (**S3: Fig.3**). The findings of this study revealed high levels of turbidity, which ranged between 1 and 4.7 NTU in standpipe water samples and between 1 and 8.1 NTU in container-stored water samples (**S4: Fig.4**). In general, the turbidity levels of both standpipe water samples and container-stored water samples were found to exceed the operational limits recommended by the South African National Standards, SANS 241 (< 1 NTU) [49].

High levels of turbidity in household drinking water imply that the storage period impacted negatively on the quality of container-stored drinking water in terms of turbidity. This could have been caused by the continuous addition of water to containers that were not cleaned over time. This practice coupled with the use of an open scoop to draw water from the container could be the source that exacerbates the increase in turbidity in stored water over time (**S2:Fig. 2**). In addition, the increase of turbidity in the standpipes could be as a result of the ageing pipes. The findings of this study corroborate the results of previous studies that attributed the deterioration of household container-stored water to the living conditions and hygiene practices in dwellings [17-19; 45; 46].

In a number of water samples, the concentrations of free chlorine residuals were found to be higher than the recommended limits (0.3 – 0.6 mg/L) [44]. This was profound in Gamalakhe. The results also showed that the target bacteria were not recorded in water samples collected from standpipes, while free chlorine residuals could not inhibit the growth of the bacteria in container-stored drinking water (**S3:Fig. 3**). This clearly confirms that high turbidity levels in container-stored water and the adverse conditions during storage in the dwellings were important sources of the microbial contamination of container-stored drinking water collected from households in Boboyi, Bomela, and Gamalakhe. High turbidity, therefore, affected the efficiency of free chlorine residual in inhibiting bacterial growth in container-stored water. Turbidity is defined as a measure of the presence of suspended particles in water that are capable of scattering light. While microorganisms are only a tiny portion of these particles (most is mud and silt), high turbidity increases the potential for transmission of infectious diseases [7; 47].

Selective media and biochemical tests were prepared and used for the identification of presumptive pathogen isolates obtained from the rural areas (Boboyi, Bomela, and Gamalakhe). Faecal coliforms, presumptive *Salmonella, Shigella* and *Vibrio* spp. were detected at high levels in container-stored water (Table 1). This means that the water did not comply with the limit for no risk of microbial infection, which is less than one faecal coliform in 100 ml of drinking water [48; 49]. Correlations were found between the high levels of turbidity in container-stored water and the incidence of presumptive bacterial pathogens. The results of this study are in agreement with previous studies, which showed strong correlations between the level of turbidity and microbial contamination of treated water [8; 13; 50]. A number of studies have also pointed out an association between increased turbidity levels and waterborne disease outbreaks [51; 52]. Gastrointestinal diseases in the elderly of Philadelphia [7] and in other Western countries has been associated with drinking-water turbidity. Our study, therefore, showed that the quality of a large number of household container-stored water samples was poor.

**Table 1:** Presence/Absence of similar bacterial pathogens detected in both household container-stored water samples and stool specimens of diarrhoeic patients after sequencing of 16S rRNA genes (n = 20 selected samples for each group, between January 2008, November 2009 and April 2015)

| Organisms | Household water | Stools HIV/AIDS +ve | Stools HIV -ve |
| --- | --- | --- | --- |
| *Escherichia coli* | P | P | A |
| *Escherichia coli* O55:H7 | P | P | A |
| *Klebsiella* spp. | P | P | A |
| *Klebsiella pneumoniae* | P | P | A |
| *Proteus mirabilis* | P | P | P |
| *Shigella boydii* | P | P | A |

+ve P = HIV-positive patient; -ve P = HIV-negative patient; P = Presence; A = Absence

Although detected when using culture-based methods, results of this study revealed that most of the target pathogenic bacteria in both household drinking water and stool specimens of HIV-positive and non-HIV patients with diarrhoea were not identified using a combination of molecular techniques. However, bacterial pathogens such as *E. coli, Vibrio, Shigella*, and *Salmonella* spp. are among the major health risks associated with water in general, and in diarrhoeal diseases in particular. Their detection by using culture-based methods alone tends to be inaccurate and also misleading in establishing an unequivocal relationship between the bacterial pathogen strains from drinking water and those from stools of HIV/AIDS individuals.

There was, therefore, a need to use robust molecular tools that could provide accurate identification pertaining to the proportion of specific pathogenic species in water samples and in stool specimens of individuals with diarrhoea. A combination of PCR targeting the 16S rRNA gene, PCR-restriction fragment length polymorphism (RFLP), and sequence analysis of 16S rRNA amplified genes resulted in the identification of similar sequences of specific bacterial pathogens from household container-stored water and those from stool specimens of HIV/AIDS-positive patients with diarrhoea. These bacterial strains included *Klebsiella* spp., *K. pneumoniae, E. coli, E. coli* O55: H7, *Proteus mirabilis*, and *Shigella boydii* (Table 1). With the exception of *Proteus mirabilis*, none of these pathogens were identified in stool specimens of non-HIV-infected individuals. These findings clearly indicated an inherent link between poor household drinking-water quality and the diarrhoeic conditions of HIV/AIDS individuals living in the Ugu District Municipality of the KwaZulu-Natal Province. These findings suggest a possibility of interspecies transmissibility between HIV/AIDS-positive patients and their household-stored drinking water.

It is well known that the function of the immune system is to protect the host against infection after exposure to bacteria, viruses, fungi, protozoa, and allergens. Protection of the host is crucial and occurs in two steps: a process of innate immunity and then adaptive T-cell recognition of self vs. non-self antigens, leading to the elimination of the organism and preservation of the internal body. If the antigen is not removed, the immune system will remain highly activated, and, in the process, burn out. The depletion of HIV-positive individuals’ immune systems predisposes this group of the population to any pathogenic organism from any source, such as contaminated drinking water. Due to their weakened immune systems, they become more susceptible to serious waterborne illnesses than persons with stronger immune systems. Contaminated drinking water, therefore, might worsen the condition of an individual who is HIV-positive. The pathogenic organism will continue to multiply within the cell for as long as there is no immune response. This clearly explains the similarity between pathogenic bacteria species in stored water samples and those in stools of HIV-positive patients who suffered from diarrhoeal disease during the study period.

*Klebsiella* spp., *K. pneumoniae, E. coli, E. coli* O55:H7, *Proteus mirabilis* and *Shigella boydii* detected in stored household drinking water samples are known to colonise the gastrointestinal tracts of humans and cause gastrointestinal diseases, ranging from simple diarrhoea to dysentery-like conditions [53-55]. There is, therefore, a higher probability for these bacterial pathogens found in household-stored water to be associated with the diarrhoeal diseases in HIV-positive patients than in HIV-negative people (Table 1). Microorganisms that appear to be harmless to the healthy individual may be fatal in immunocompromised persons such as HIV/AIDS individuals and the elderly. In HIV-negative individuals, normal floras are not easily depleted and can overcome the opportunistic microorganisms. Therefore, their strong immune system protects them by providing a stronger and more effective immune response after exposure to most of the pathogenic bacteria found in stored water, except *Proteus* spp. This study suggests that only these organisms were associated with the diarrhoeal disease in HIV-negative individuals (Table 1).

Poor water quality remains a health hazard for HIV/AIDS individuals in the Ugu District Municipality and in other rural areas of South Africa. These findings support studies conducted by previous investigators in the Limpopo [11; 13] and Eastern Cape Provinces of South Africa, who also linked unsafe drinking water to diarrhoeic conditions of HIV/AIDS individuals. Pathogenic bacteria such *Salmonella enterica* serovar Enteritidis *(S.* Enteritidis), *Salmonella Typhimurium, E. coli*, and *Shigella dysenteriae* found in Limpopo Province drinking water were also detected in stool specimens of HIV-negative individuals with and without diarrhoea, but there was a higher occurrence in HIV/AIDS individuals [11]. In the Eastern Cape, *E. coli* O157: H7 detected in the stool specimens of HIV/AIDS patients visiting Frere Hospital for a diarrhoeal disease was linked to drinking-water sources used by the communities [55]. All these studies provide clear evidence that drinking-water sources used by rural communities of the poorest provinces in the country are at high risk of increasing diarrhoeal diseases in immunocompromised people such as HIV/AIDS individuals.

## 5. Conclusion and recommendation

The findings of this study indicated a higher possibility for *Klebsiella* spp., *K. pneumoniae, E. coli, E. coli* O55: H7, *Proteus mirabilis* and *Shigella boydii* in the stored household drinking water of Boboyi, Bomela and Gamalakhe to be associated with the diarrhoeic conditions of HIV/AIDS patients visiting the Murchison and Port Shepstone hospitals in the Ugu District Municipality of KwaZulu-Natal. The poor quality of water stored in households in this district had a major impact on the health of immunocompromised individuals. The provision of safe water is therefore imperative to safeguard public health in general and that of HIV-infected individuals in particular. This study recommends the urgent implementation of sanitary conditions and practices for the storage of drinking water in households in order to minimise the number of pathogens entering the household container-stored water.

## 6. Acknowledgement

We thank the National Research Foundation of South Africa and the Tshwane University of Technology for funding this study.

## Supporting documents

S1: Fig. 1

S2: Fig. 2

S3: Fig. 3

S4: Fig. 4

---

**S1 Fig. 2:**
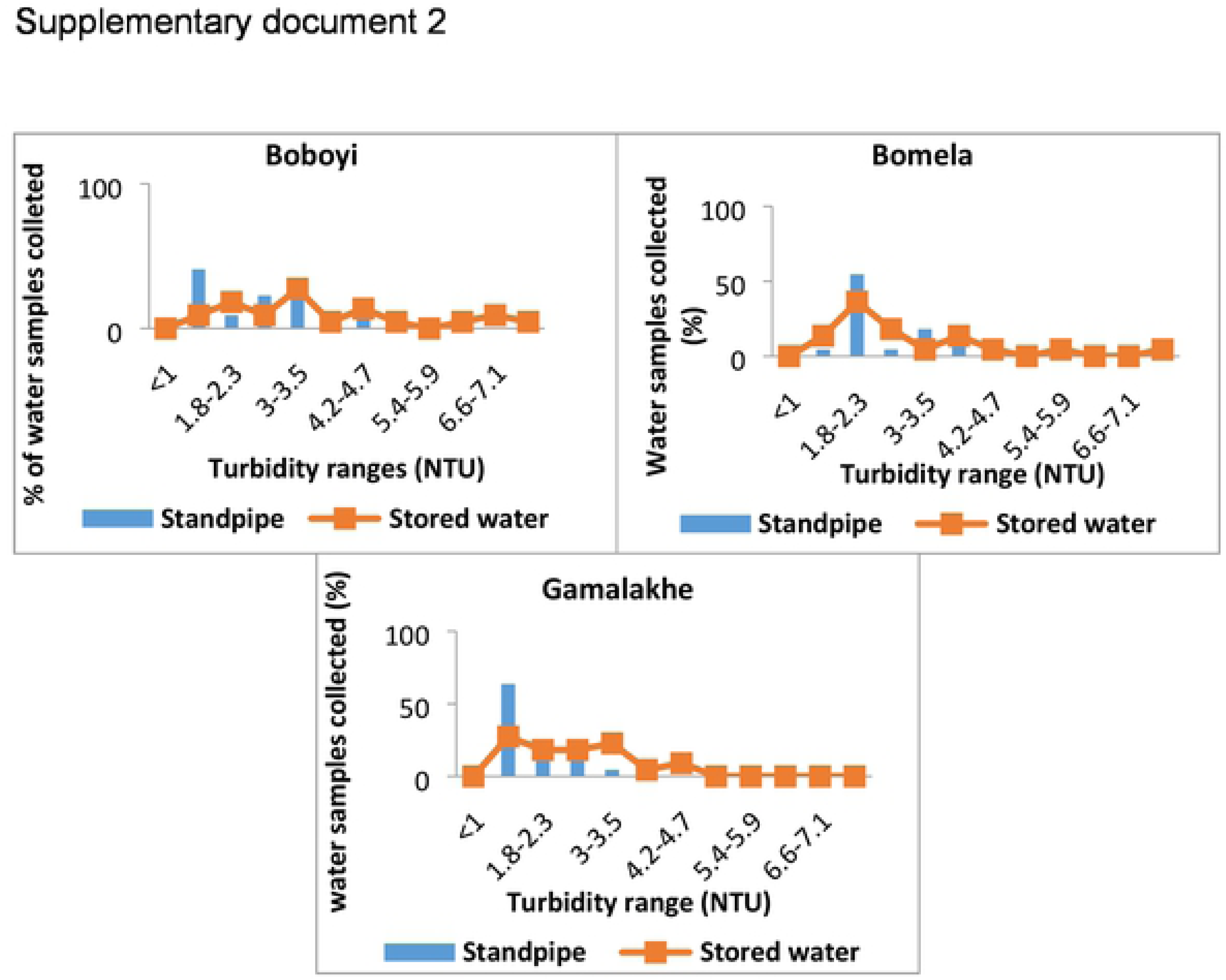
The average turbidity values (NTU) in standpipes and stored household container water samples collected from Boboyi, Bomela and Gamalakhe

## References

1. Momba, M.N.B., Tyafa, Z., Makala, N., Brouckaert, B.M. & Obi, C.L. 2006b. Safe drinking water still a dream in rural areas of South Africa. Case Study: The Eastern Cape Province. Water SA, 32(5): 715–720.

2. Republic Of South Africa. 1996. Constitution of the Republic of South Africa, 1996 (Act No. 108 of 1996). Government Gazette, 378 (147):176–178.

3. Duse, A.G., Da Silver, M.P. & Zietsman, I. 2003. Coping with hygiene in South Africa, a water scare country. International Journal of Environmental Health Research, 13: 95–105.

4. Kasrils, R. 2004. A Decade of Delivery. Minister of Water Affairs and Forestry, Pretoria, South Africa.

5. Department Of Water Affairs & Forestry (DWAF). 2008. Water and Sanitation Coverage in South Africa. Department of Water Affairs and Forestry, Pretoria, South Africa.

6. WHO (World Health Organization) & UNICEF 2015. Progress on Sanitation and Drinking water: 2015 Update and MDG Assessment. ISBN 978-92-4-150329-7. UNICEF and WHO, New York, USA.

7. Schwartz, J., Levin, R. & Goldstein, R. 2000. Drinking water turbidity and gastrointestinal illness in the elderly of Philadelphia. Journal of Epidemiology and Community Health, 54: 45–51.

8. Momba, M.N.B & Kaleni, P. 2002. Regrowth and survival of indicator microorganisms on the surfaces of household containers used for the storage of drinking water in rural communities of South Africa. Water Research, 36: 3023–3028.

9. Obi, C.L. & Bessong, P.O. 2002. Diarrhoegenic bacterial pathogens in HIV-positive patients with diarrhoea in rural communities of Limpopo Province, South Africa. Journal of Health Population and Nutrition, 20: 230–234.

10. Momba, M. N. B., Tyafa, Z & Makala., N. 2004a. Rural water treatment plants fail to provide potasble water to their consumers: Alice water treatment plant in the Eastern Cape Province of South Africa. South African Journal of Science, 100: 307–310.

11. Obi, C.L, Onabolu, B., Momba, M.N.B., Igumbor, J.O., Ramalivahna, J., Bessong, P.O., Van Rensburg, E.J., Lukoto, M., Green, E. & Mulaudzi. T.B. 2006. The interesting cross-paths of HIV/AIDS and water in Southern Africa with special reference to South Africa. Water SA, 32: 56–78.

12. Obi, C.L., Ramalivhana, J., Momba, M.N.B. & Igumbor, J. 2007a. Scope and frequency of enteric bacterial pathogens isolated from HIV/AIDS patients and their household drinking water in Limpopo Province. Water SA, 33: 1–10.

13. Momba, M.N.B., Tyafa, Z & Makala, N. 2003. Rural water treatment plants fail to provide potable water to their consumers: Alice water treatment plant in the Eastern Cape Province of South Africa. South African Journal of Science, 100: 307–310.

14. Bailey, I.W. & Archer, L. 2004. The impact of introducing treated water on aspects of community health in a rural community in KwaZulu-Natal. Water Science and Technology, 50: 105–110.

15. Seino, K., Takano, T., Quang, N.K.L., Wanatabe, M., Inose, T. & Nakamura, K. 2008. Bacterial quality of drinking water stored in containers by boat households in Hue City, Vietnam. Environmental Health and Preventive Medicine, 13: 198–206.

16. Maraj, S., Rodda, N., Jackson, S., Buckley, C. & Macleod, N. 2010. Microbial deterioration of stored water for users supplied by stand-pipes and ground-tanks in a peri-urban community. Water South Africa, 32.

17. Jagals, P., Bokako, T. & Grabow, W. 1999. Changing consumer water-use patterns and their effect on microbiological water quality as a result of an engineering intervention. Water South Africa, 25: 297–300.

18. Trevett, A. 2003. The public health significance of drinking-water quality deterioration in rural Honduran communities. PhD Thesis, Silsoe College, Cranfield University, Bedfordshire, UK.

19. Gundry, S., Wright, J. & Conroy, R. 2004. A systematic review of the health outcomes related to household water quality in developing countries. Journal of Water and Health, 2: 1–13.

20. Trevett, A.F., Carter, R.C. & Tyrrel, S.F. 2005. The importance of domestic water quality management in the context of faecal-oral disease transmission. Journal of Water and Health. 3(3): 221–228.

21. Tambekar, D.H. & Banginwar, Y.S. 2004. Studies on intervention for control of water borne diseases: Promoting personal and domestic hygiene in hotels/restaurant’s owner and workers. Journal Comparative Toxicology and Physiology, 1: 267–276.

22. Tambekar, D.H., Gulhane, S.R., Jaisingkar, R.S., Wangikar, M.S., Banginwar, Y.S & Mogarekar, M.R. 2008. Household water management: A systematic study of bacteriological contamination between source and point of use. Journal of Agriculture and Environmental Science, 3: 241–246.

23. Sobsey, M.D., Handzel, T. & Venczel, L. 2003. Chlorination and safe storage of household drinking water in developing countries to reduce waterborne disease. Journal of Water Science and Technology, 47: 221–228.

24. Hunter, P.R., Waite, M. & Ronchi, E. 2002. Drinking Water and Infectious Disease: Establishing the Links. IWA Publishing, London, UK.

25. Ashbolt, N. J. 2004. Microbial contamination of drinking water and disease outcomes in developing regions. Toxicology, 198: 229–238.

26. WHO (World Health Organization). 2003. Emerging Issues in Water and Infectious Disease Series. J. Bartram, J. Cotruvo, M. Exner, C. Fricker, A. Glasmacher (Eds.), Heterotrophic Plate Counts and Drinking-Water Safety: The Significance of HPCs for Water Quality and Human Health. Published on behalf of WHO by IWA Publishers, London, UK.

27. HSRC (Human Sciences Research Council). 2002. Nelson Mandela HSRC study of HIV/AIDS. ISBN 0-7069-2007-9.

28. Lakshminarayanan, S & Ramakrishnan, J. 2015. Diarrheal diseases among children in India: Current scenario and future perspectives. Journal of Natural Science, 6(1): 24–28.

29. WHO (World Health Organization) 2002c. WHO Statistical Information System (WHOSIS). Geneva, Switzerland.

30. Stats SA. 2015a. Millennium Development Goals: Country report 2015. Millennium Development Goals: Country Report 2015/Statistics South Africa. Statistics South Africa, Pretoria, South Africa. ISBN 978-0-621-43861-1.

31. Hayes, C.E., Elliot, E., Krales, E. & Goulda, D. 2003. Food and water safety for persons infected with human immunodeficiency virus. Clinical Infectious Disease, 36: 106–109.

32. Ashton, P.J. 2000. The potential of HIV/AIDS for the Thukela Water Project. Contract Report to the Department of Water Affair and Forestry, Pretoria, South Africa.

33. Obi, C. L., Ramalivhana, J., Momba, M.N.B., Onabolu, B., Igumbor, J.O., Lukoto, M., Mulaudzi, T.B., Bessong, P.O., Jansen Van Rensburg, E.L., Green, E. & Ndou, S. 2007b. Antibiotic resistance profiles and relatedness of enteric bacterial pathogens isolated from HIV/AIDS patients with and without diarrhoea and their household drinking water in rural communities in Limpopo Province South Africa. African Journal of Biotechnology, 6: 1035–1047.

34. Abong’o, B.O., Momba, M.B.N., Makalate, V. K. & Mwambakana, J.N. 2008. Prevalence of *Escherichia coli* O157:H7 among diarrhoeic HIV/AIDS patients in the Eastern Cape Province-South Africa. Journal of Biological Science, 11: 1066–1075.

35. Stats SA. 2009. Stats SA, Mid-year population estimates, South Africa, 2009 (Statistical release P0302).

36. APHA, 1998. Standard Methods for the Examination of Water and Wastewater. Joint publication of the American Public Health Association (APHA), American Water Works Association (AWWA) and Water Environment Federation (WEF). www.apha.org, www.awwa.org

37. South African Bureau Of Standards (SABS). 2001. Water quality-detection and enumeration of *Vibrio cholerae*. Standard Methods, 1st edn. SABS-SB 1315, Pretoria.

38. APHA, 2005. Standard Methods for the Examination of Water and Wastewater. Joint publication of the American Public Health Association (APHA), American Water Works Association (AWWA) and Water Environment Federation (WEF), www.apha.org, www.awwa.org.

39. Lane, D.J. 1991. 16S/23S rRNA sequencing. In: Stackebrandt, E. and Goodfellow, M. (Eds.), Nucleic Acid Techniques in Bacterial Systematics. John Wiley and Sons: 115-175.

40. Heyndrickx, M., De Vos, P. & De Ley, J. 1991. Fermentation characteristics of *Clostridium pasteurianum* LMG 3285 grown on glucose and mannitol. Journal of Applied Bacteriology, 70: 52–58.

41. Hall, T.A. 1999. BioEdit: a user-friendly biological sequence alignment editor and analysis program for Windows 95/98/NT. Nucleic Acids, 41: 95–98.

42. Lechevallier, M.W., Abbaszadegan, M. & Camper., A.K. 1999. Committee Report: emerging pathogens-bacteria. Journal of American Water Work Association, 91: 101–109.

43. Momba, M.N.B., Cloete, T.E., Venter, S.N. & Kfir, R. 1999. Examination of the behaviour of *Escherichia coli* in biofilms established in laboratory scale units receiving chlorinated and monochlorinated water. Water Research, 33, 2937–2940.

44. DWAF, DoH & WRC, 1998. Quality of Domestic Water Supplies, Volume 1: Assessment Guide, 2nd Edn. Water Research Commission Report No. TT 101/98. Department of Water Affairs and Forestry, Department of Health, Water Research Commission (WRC), Pretoria, South Africa. http://www.dwaf.gov.za/IWQS/AssessmentGuides/AssessmentGuide/Assessmentguide.pdf

45. Oswald, W.E., Lescano, A.G., Bern, C., Calderon, M.M., Cabrera, L. & Gilman, R.H. 2007. Faecal contamination of drinking water within peri-urban households, Lima, Peru. Journal of Tropical Medicine and Hygiene, 77: 699–704.

46. Rufener, S., Mausezah, D., Mosler, H. & Weingartner, J. 2010. Quality of drinking-water at source and point of consumption–drinking cup as a high potential recontamination risk: A field study in Bolivia. Journal of Health, Population and Nutrition, 28(1): 34–41.

47. Department Of Water Affairs & Forestry (DWAF) 1996. South African Water Quality Guidelines – Volume 1: Domestic Water Use. Department of Water Affairs and Forestry, Pretoria, South Africa, 86–87.

48. DWAF, DoH & WRC, 1998. Quality of Domestic Water Supplies, Volume 1: Assessment Guide, 2nd Edn. Water Research Commission Report No. TT 101/98. Department of Water Affairs and Forestry, Department of Health, Water Research Commission (WRC), Pretoria, South Africa. http://www.dwaf.gov.za/IWQS/AssessmentGuides/AssessmentGuide/Assessmentguide.pdf

49. SANS 241. 2006. South African National Standard: Drinking Water Specification, Edition 6.1. South African National Standard 241: 2006. South African Bureau of Standards, Pretoria, South Africa.

50. Lechevallier, M.W. & Norton, W. D. 2000. Treatments to address source water concerns: Protozoa. Drinking water turbidity and for gastrointestinal illness in elderly of Philadelphia balancing chemical and microbial risks. Epidemiology, 8: 615–620.

51. Kent, J.P., Greenspan, J.R. & Herndon, J.L. 1988. Epidemic giardiasis caused by a contaminated public water supply. American Journal of Public Health, 78: 139–43.

52. Mackenzie, W. R., Kazmierczak, J.J. & Davis, J.P. 1994. A massive outbreak in Milwaukee cryptosporidium infection transmitted through the public water supply. New England. Journal of Medicine, 331: 161–167.

53. Janda, J.M., & Abbott, S.L. 2010. The Genus Aeromonas: Taxonomy, Pathogenicity, and Infection. Clinical Microbiology Reviews, 23: 35–73. http://dx.doi.org/10.1128/CMR.00039-09.

54. Martínez, J., Martínez, L., Rosenblueth, M., Silva, J. & Martínez-Romero, E. 2004. How are gene sequence analyses modifying bacterial taxonomy? The case of *Klebsiella -* Review Article. International Microbiology, 7: 261–268.

55. Von Seidlein, L., Kim, D.R., Ali, M., Lee, H. & Wang, X.Y. 2006. A multicentre study of *Shigella* diarrhoea in six Asian countries: Disease burden, clinical manifestations, and microbiology. PLoS Medicine, 3: 351–353.

